# Heterogeneity in clone dynamics within and adjacent to intestinal tumours identified by Dre-mediated lineage tracing

**DOI:** 10.1101/2020.05.13.094284

**Authors:** Ann-Sofie Thorsen, Doran Khamis, Richard Kemp, Mathilde Colombé, Filipe C. Lourenço, Edward Morrissey, Douglas Winton

**Affiliations:** Cancer Research-UK Cambridge Institute, Li Ka Shing Centre, Robinson Way, Cambridge, CB2 0RE, UK; University of Oxford, Center for computational biology, Weatherall Institute of Molecular Medicine, University of Oxford, John Radcliffe Hospital, Oxford OX3 9DS, UK

**Keywords:** Lineage tracing, Cancer, Intestine, Epithelial

## Abstract

Somatic models of tissue pathology commonly utilise induction of gene specific mutations in mice mediated by spatiotemporal regulation of Cre recombinase. Subsequent investigation of the onset and development of disease can be limited by the inability to track changing cellular behaviours over time. Here a lineage tracing approach based on ligand dependent activation of Dre recombinase that can be employed independently of Cre is described. The clonal biology of intestinal epithelium following Cre-mediated stabilisation of ß-catenin reveals that within tumours many new clones rapidly become extinct. Surviving clones show accelerated population of tumour glands compared to normal intestinal crypts but in a non-uniform manner indicating that intra-tumour glands follow heterogeneous dynamics. In tumour adjacent epithelium clone sizes are smaller than in the background epithelium as a whole. This suggests a zone of around 5 crypt diameters within which clone expansion is inhibited by tumours and that may facilitate their growth.

## Introduction

Mouse models of pathology in which phenotypes are somatically induced by the directed expression of recombinases have become a ubiquitous tool across all branches of the medical sciences. Currently there are over 4000 mouse lines engineered for this purpose (EUCOMM, 2019; Jax.org, 2019). Activation of recombination in adult tissues is highly efficient and can result in altered cellular behaviours that can change over time due to adaptation, cellular exhaustion or progression and consequently display different phenotypes that may reflect different disease settings. Examples include arteriosclerosis (Ishibashi et al., 1993), diabetes (Zhang et al., 1994), inflammation (Stremmel et al., 2017), Alzheimer’s (Matsuda et al., 2008), Parkinson’s disease (Choi et al., 2017) and cancer (Cheung et al., 2009).

An important aspect of the phenotypic characterisation of affected tissues following recombination often includes lineage tracing in which the origins and fate of individual cells and their descendants are followed over time using acquired expression of a cell autonomous reporter gene. Problematic in such lineage tracing experiments is that reporter expression is often dependent on the activity of the same recombinases acting to induce the phenotype of interest. This is a major limitation as the level and timing of recombination required for lineage tracing may be very different from that needed to induce the phenotype although re-switchable cassettes can sometimes be employed (Schepers et al., 2012).

The most widely used DNA recombinases in *in vivo* mouse models are Cre and Flp (Sadowski, 1995; Sternberg and Hamilton, 1981). Historically, most cancer associated conditional alleles contain pairs of loxP sites for Cre driven recombination: e.g. *Tp53*^flox2-10^ (Marino et al., 2000), *Apc*^flox^ (Cheung et al., 2009) and *Ctnnb1*^flox(ex3)^ (Harada et al., 1999). Some frt alleles for Flp driven recombination are also available: e.g. *Kras^fsfG12D^* (Young et al., 2011). To employ recombinases sequentially requires independent spatiotemporal control of the activity of different DNA-recombinases that can only occur if they are expressed under different promoters and/or activated by different ligands.

The novel Dre DNA recombinase, discovered in a screen for Cre-like enzymes (Sauer and McDermott, 2004), recognises 32 bp rox sites. The Dre/rox system does not cross-react with the Cre/loxP system (Anastassiadis et al., 2009). Dre has been used in conjunction with Cre and Flp to identify cell populations defined by differential promoter activity (Hermann et al., 2014; Madisen et al., 2015; Plummer et al., 2015; Sajgo et al., 2014). An inducible and functional Dre^Pr^ fusion protein (Dre fused to the human progesterone receptor and activatable by the synthetic analogue Ru486) has been described (Anastassiadis et al., 2009) but has been used *in vivo* only in a single Zebrafish study (Park and Leach, 2013). Here we employ Dre^Pr^ for somatic studies in adult mice and demonstrate that it can be used in combination with tamoxifen inducible Cre^Ert^ alleles to initiate lineage tracing at any time following the activation of a Cre-mediated phenotype. The method is applied to determine the altered fate and clone dynamics of stem cell populations within and adjacent to intestinal tumours induced by stabilisation of β-catenin.

## Results

### RDre^Pr^ mice have widespread Dre expression

The RDre and RDre^Pr^ animals were created by homologous recombination of targeting vectors into the *R26* locus in mouse embryonic stem cells as described before (Vooijs, et al., 2001) (Figure S1A). A rox-STOP-rox (rsr) tdTomato (tdTom) reporter line (*R26*^rsrtdTom^) was generated by germline deletion of the loxP-STOP-loxP (lsl) cassette from the *R26*^rsrlslTdTom^ allele (The Jackson Laboratory stock no. 021876). To investigate Dre activity in different tissues, RDre and RDre^Pr^ animals were crossed to these *R26^rsrtdTom^* reporter mice to generate compound *RDre;R26^rsrtdTom^* and *RDre^Pr^;R26^rsrtdTom^* here referred to as RDre;RtdTom^rsr^ and RDre^Pr^;RtdTom^rsr^, respectively.

First, adult RDre;RtdTom^rsr^ mice were analysed and the widespread activity of Dre across many tissues was confirmed by IHC for tdTom expression (Figures S1B and S2). However, cells in the outer most layer of the epidermis and bone cartilage did not express any tdTom at time of analysis (Figures S1B and S2). Next, to investigate Dre^Pr^ activity after Ru486 exposure, RDre^Pr^;RtdTom^rsr^ animals had 1, 2 or 3, 90-day slow release pellets containing 10mg/pellet Ru486 implanted sub-cutaneously. At 75 days post implantation animals were sacrificed, and tissues from different germ-layers were collected for tdTom expression analysis. Importantly, the Dre^Pr^ fusion mice showed a complete absence of tdTom expression in all tissues in uninduced mice (Figures S1B and S2). In contrast, following induction with Ru486 sporadic tdTom expression was observed in many tissues. Epithelial cells and/or clones expressing tdTom were observed in: endoderm derived intestine, stomach, liver, pancreas; mesoderm derived spleen and kidney and ectoderm derived skin (Figure S1B). Furthermore, quantitative flow cytometry confirmed that intestinal epithelial cells were activated by Ru486 pellets in a dose responsive manner (Figure S3A-C). Of note, not all tissue types analysed expressed tdTom after Ru486 exposure including lung, heart, bone and tongue (Figure S2). The observation that these tissues did show widespread recombination in RDre;RtdTom^rsr^ mice suggests transcriptional silencing at the *ROSA* locus in these tissues in adult mice or that Ru486 does not reach all tissues, as described before for Cre recombiase and Tamoxifen (Sinha and Lowell, 2017; Vooijs et al., 2001). These results show that Dre^Pr^ has no background activity and is activated by Ru486 in various tissues, including the intestinal epithelium.

### Dre^Pr^ can be clonally induced in epithelial cells

Next, the ability of Dre^Pr^ to operate as a lineage tracing tool in epithelial tissues was investigated. Lineage tracing is commonly carried out by pulse chase experiments (Ghosh et al., 2011; Giroux et al., 2017; Van Keymeulen et al., 2011; Papafotiou et al., 2016; Snippert et al., 2010; Vermeulen et al., 2013). To investigate the activity of Dre^Pr^ after a pulse of inducer, Ru486 was administered to RDre^Pr^;RtdTom^rsr^ mice by intra peritoneal (i.p.) injection at doses of 50 or 80 mg/kg (single injections) or 240 mg/kg (80mg/kg administered on three consecutive days). After 14 days animals were culled and the bladder, trachea, oesophagus and caecum were analysed by fluorescent microscopy (Figure 1A-D). This revealed that tdTom positive clones could be observed in all tissues (Figure 1A-D). Additionally, clones were also observed in the intestine (Figure 1E-G) and the number of clones observed in the small intestine and colon 14 days post induction increased in a dose responsive fashion in both (Figure 1H,I). The number of clones/cm in the small intestine was ~100 vs ~300 following a single or three dose(s) of 80 mg/kg Ru486, respectively, and indicated a near linear accumulation of signal (Figure 1H,I). Twenty-four hours after Ru486 administration, single tdTom+ cells could be observed in the bottom of intestinal crypts (Figure 1J). Over time, tdTom could be observed in whole crypts and villi after Dre^Pr^ induction both in the small intestine and colon (Figure 1K-O). Such, fully clonal crypt-villus clones expressed both goblet, Paneth, Tuft and enteroendocrine cells (Figure 1K-O), underlining that Dre^Pr^ was activated in single clonal intestinal stem cells, and that these cells can give rise to differentiated daughter cells.

**Figure 1:**
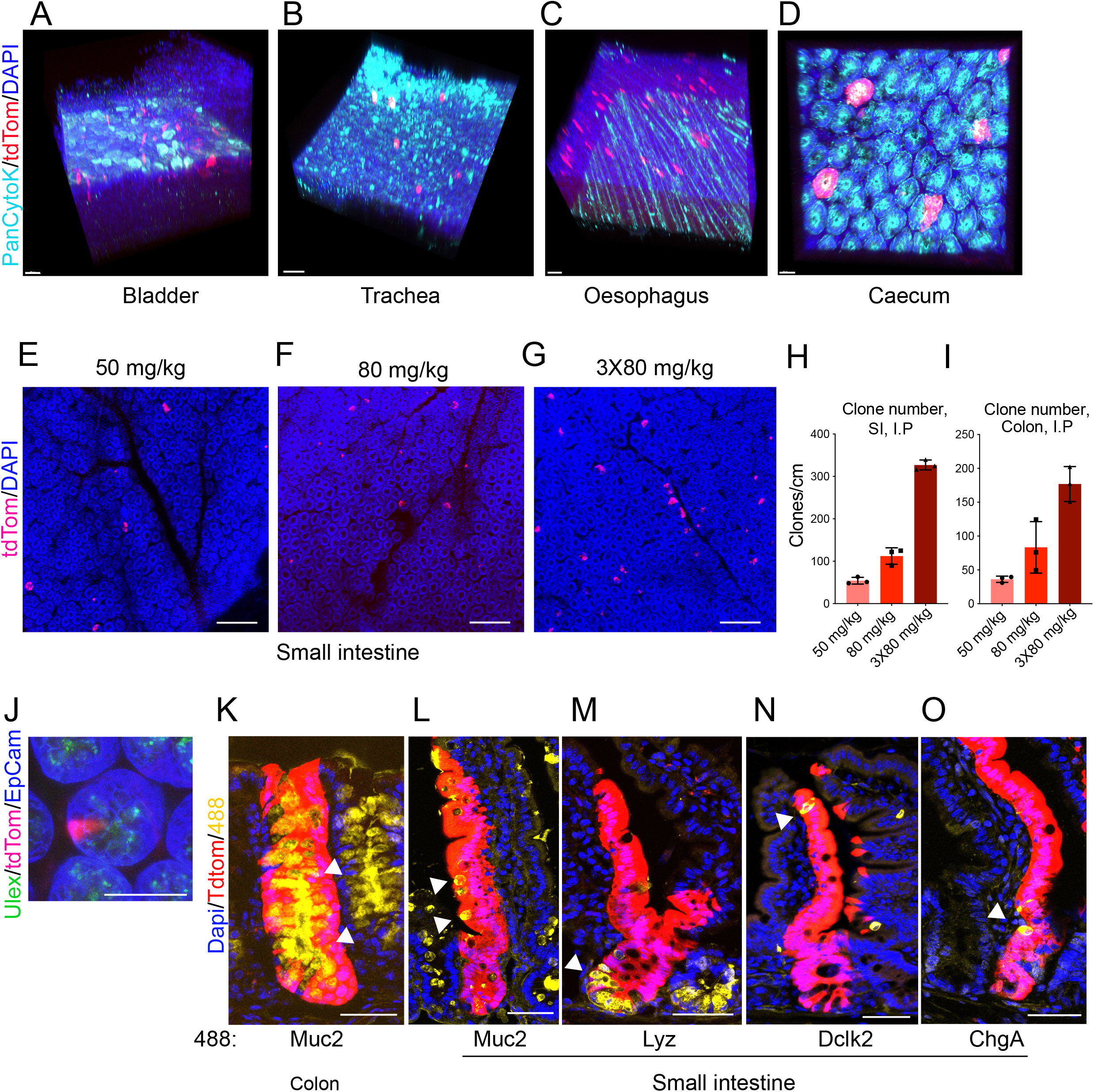
Clonal recombination induced by Dre^Pr^. **(A-D)** Expression of tdTomato in tissues shown from RDre^Pr^;RtdTom^rsr^ animals 14 days following i.p. injection of 240 mg Ru486. **(E-G)** Confocal images showing expression of tdTomato in the crypts of small intestinal wholemounts of RDre^Pr^;RtdTom^rsr^ animals 14 days following i.p. injection of Ru486 at the doses shown **(H, I)** Bar-graph showing quantification of the number of tdTom+ crypt clones/cm in small intestinal (H) or colon tissue (I), (N=3; mean +/- SD). **(J)** small intestinal crypt from a RDre^Pr^;RtdTom^rsr^ animal 24 hours after receiving 50 mg/kg Ru486 by i.p. injection showing a single tdTomato+ cell not expressing Paneth cell marker (Ulex lectin). **(K-O)** Confocal images of tdTomato+ clones from the same animals as for E-I. Colon section visualised for Goblet cells (Muc2) (K). Small intestinal sections (L-O) visualised for Goblet cells (Muc2) (L), visualised for Paneth cells (Lysozyme) (M), visualised for tuft cells (Dlck2) (N) and visualised for enteroendocrine cells (Chromogranin A) (O). All scalebars = 50μm.

For clonal lineage tracing experiments to be informative the frequency of clone induction has to be low enough to avoid clonal collisions over the time course of the experiment but high enough to permit quantitation within a defined region of the tissue. Here we find that 50 mg/kg Ru486 was the optimal pulsing dose in lineage trace experiments as this dose appeared to hit <1 stem cell per crypt at early time points (Figure 1J) and induced an appropriate sparsity of clones at later timepoints (Figure 1E, H,I).

### Lineage tracing and quantitative inference of intestinal stem cell dynamics using Dre^Pr^

The efficacy of Dre^Pr^ for lineage tracing was investigated in mouse intestine by performing a direct quantitative comparison of epithelial clone size distributions to those obtained in a previously employed Cre-based model (Kemp, 2004; Vermeulen et al., 2013). Clones were induced in AhCre^Ert^;RtdTom^lsl^ mice by a single induction dose (40 mg/kg ß-naphthoflavone and 0.15 mg Tamoxifen) and RDre^Pr^;RtdTom^rsr^ mice received a single dose of 50 mg/kg Ru486. Small intestine and colon from both strains were analysed at 4, 7, 10, 14 and 21 days post induction and fluorescence microscopy of whole mounted tissues was employed to score the relative sizes of tdTom+ clones in intestinal crypts (Figure 2 A,B). The average clone sizes and changes in the clone size distribution with time in the small intestine and colon were found to be remarkably similar in the two models (Figure 2C-H).

**Figure 2:**
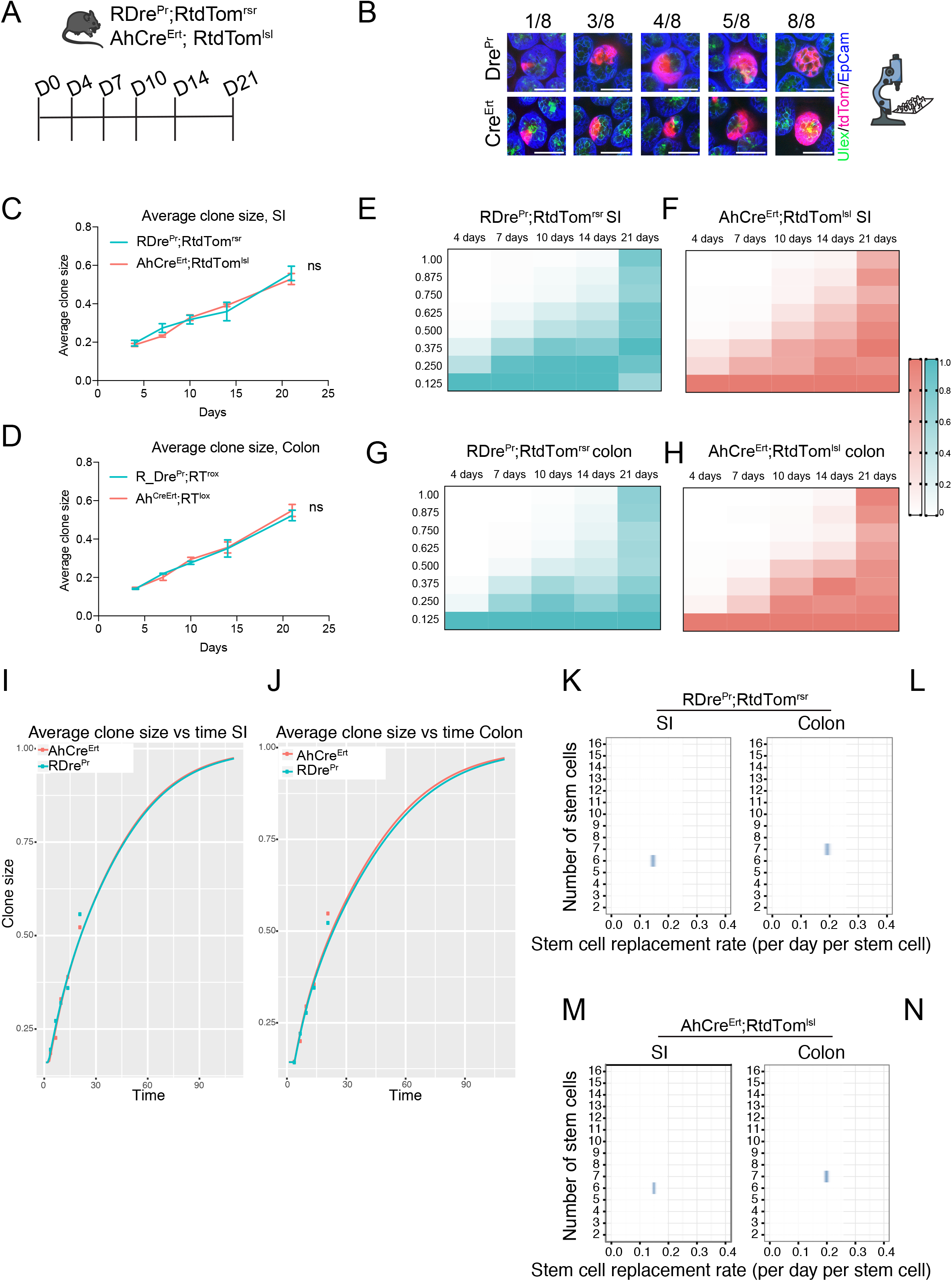
Stem cell dynamics inferred from intestinal clone size distributions in RDre^Pr^;RtdTom^rsr^ mice. (**A**) Schematic of experimental set-up. RDre^Pr^;RtdTom^rsr^ and AhCre^Ert^;RtdTom^lsl^ animals were administered a single dose of 50mg/kg Ru486 or 40mg/kg β-naphthoflavone (BNF) plus 0.15mg Tamoxifen (Tam), respectively. (**B**) Confocal microscopy images of small intestinal clones demonstrating segmental scoring in eighths of clone sizes (viewed from crypt base) from RDre^Pr^;RtdTom^rsr^ and AhCre^Ert^;RtdTom^lsl^ animals. (Scalebar= 50μm) (**C,D**) Average clone sizes with time in small intestine (SI) (C) and colon (D) derived from RDre^Pr^;RtdTom^rsr^ and AhCre^Ert^;RtdTom^lsl^ animals. (Mean+/- SD) (N=3). (**E-H**) Heatmap representing colour-coded clone size prevalence over time during lineage tracing in RDre^Pr^;RtdTom^rsr^ SI (E) and colon (G) or AhCre^Ert^;RtdTom^lsl^ SI (F) and colon (H). Darkest colour corresponds to most prevalent clone sizes at a given timepoint. (**I-J**) Mathematical modelling (line) showing predicted average clone sizes from day 0-100 in small intestine (SI) (I) and colon (J) based on the data shown in C,D (points). (**K-N**) Inferred stem cell number per crypt (y-axis) and stem cell replacement rate per stem cell per day (x-axis) in SI (K) and colon (L) of RDre^Pr^;RtdTom^rsr^ animals and SI (M) and colon (N) AhCre^Ert^;RtdTom^lsl^ animals.

Changes in clone size distributions with time arise from a neutral drift pattern of stem cell renewal that is characterised in each crypt as a 1D random walk on a ring of N stem cells each with a daily replacement rate λ (Kozar et al., 2013; Lopez-Garcia et al., 2010). For RDre^Pr^;RtdTom^rsr^ and AhCre^Ert^;RtdTom^lsl^ animals (Figure 2I-N) the model predicted average clone sizes over 100 days based on the lineage tracing data (Figure 2I-J). The analysis inferred N to be 6 and 7 in the small intestine and colon, respectively, in both the RDre^Pr^;RtdTom^rsr^ and AhCre^Ert^;RtdTom^lsl^ models (Figure 2K-L). Furthermore, λ was estimated to be 0.15 vs 0.15 in the small intestine and to 0.19 vs 0.20 in the colon of RDre^Pr^;RtdTom^rsr^ and AhCre^Ert^;RtdTom^lsl^ animals, respectively (Figure 2K-L). Importantly the average clone sizes presented here and the inferred values for N and λ were similar to those previously reported using alternative Cre recombination models or orthogonal approaches (Kozar et al., 2013; Vermeulen et al., 2013). Taken together, these observations show that the epithelial behaviours of intestinal stem cells are not perturbed by either Dre^Pr^ expression or the treatment with Ru486 indicating that Dre^Pr^ traced stem cells have a non-biased neutral behaviour.

### Dre^Pr^ can trace single cell derived clones in intestinal tumours

Performing lineage tracing within overt or developing Cre-mediated pathologies requires a Cre-independent mechanism for reporter activation. To test the ability of Dre^Pr^ to act in this way a Cre-induced intestinal tumour model based on stabilisation of ß-catenin was employed. In *Ctnnb1*^flox(ex3)^ mice removal of floxed exon3 is sufficient to induce development of intestinal tumours (Harada et al., 1999). Here, AhCre^Ert^ was utilized to induce *Ctnnb1* recombination and tumour initiation and Dre^Pr^ was utilized to trace clones derived from single cells within nascent and established tumours.

To determine the number of traceable cells per tumour *Ctnnb1* was first recombined in the intestinal epithelium of AhCre^Ert^;*Ctnnb1*^lox(ex3)^;RDre^Pr^;RtdTom^rsr^ animals by activating AhCre^Ert^ with β-naphthoflavone and Tamoxifen. Importantly, these drugs did not induce tdTom expression (Figure 3A-B and Figure S4). At 21days post *Ctnnb1* recombination lineage tracing in tumours was initiated by RDre^Pr^ induction and mice were then aged until maximum tumour burden, 13 days (N=2), 17 days (N=1) and 19 days (N=1) days post Ru486 induction (Figure 3C). This protocol produced intestinal tumours expressing stabilised β-catenin that contained tdTom+ clones (Figure 3D-G).

**Figure 3:**
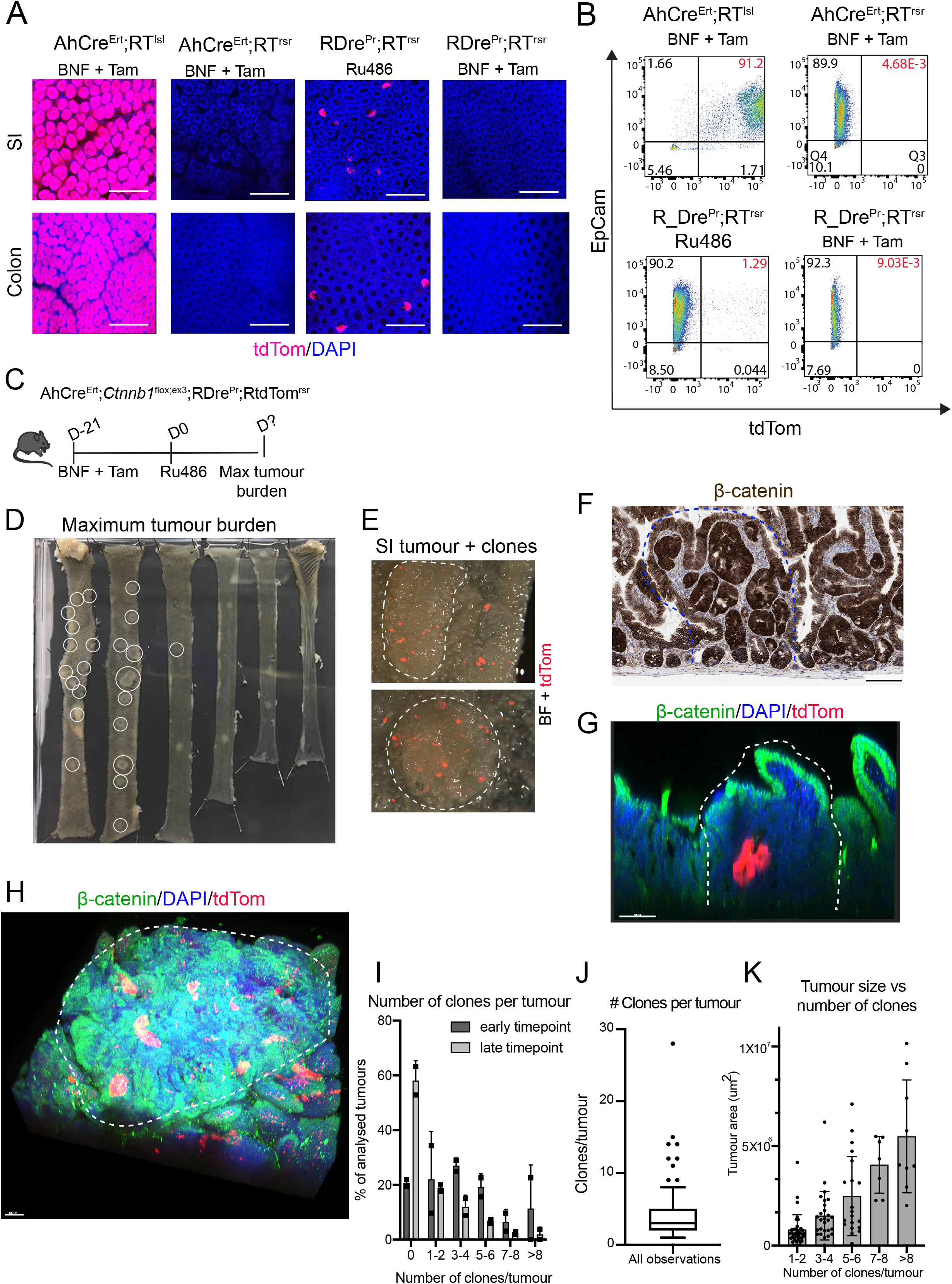
Activation of Dre^Pr^ allows lineage tracing within intestinal tumours initiated by Cre-mediated stabilisation of ß-catenin. (**A-B**) ß-naphthoflavone (BNF) and Tamoxifen (Tam) induces recombination of loxP but not rox sites in AhCre^Ert^ animals and not at rox sites in RDre^Pr^ animals. (A) Confocal microscopy images of small intestine and colon from AhCre^Ert^;RtdTom^lsl^, AhCre^Ert^;RtdTom^rsr^ and RDre^Pr^;RtdTom^rsr^ animals treated as shown. Scale bar=200μm (B) Flow cytometry quantification of tdTom+ cells in proximal small intestine from above animals. Cells were stained with EpCam (Alexa 647) and DAPI. (**C**) Schematic representation of experimental set-up. Tissue was collected at humane endpoint. (**D**) Whole-mount of intestine at maximum tumour burden. Macroscopic tumours are circled. (**E**) Dissection microscopy picture of whole-mounted small intestinal (SI) tumours with tdTom+ clones. Overlay of the brightfield (BF) and 555nm channel (tdTom). (**F**) Section showing intestinal tumour (blue line) with IHC for ß-catenin. Scalebar = 100μm (**G**) Cross-section of a tumour (white line), containing a tdTom+ clone. Scalebar = 100μm. (**H**) Representative confocal microscopy images of multiple tdTom+ clones in tumour stained for β-catenin (white line). Scalebar = 100μm. (**I**) Bar-graph summarising the distribution of number of tdTom+ clones (0 to >8) in tumours from mice culled 13 days (early) or 17-19 days (late) post Ru486 induction (N=62 and 121 respectively). Each datapoint shows mean number of tumours per mouse. (**J**) Box-plot displaying the average number of clones per tumour (of all tumours containing clones), N=100, data outside the 10th and 90th percentile is displayed as single datapoints. (**K**) Bar-graph showing the number of tdTom+ clones (1 to >8, binned by 2) in all tumours based on tumour area (μm^2^). Each datapoint is a tumour, N=100, error bars = SD.

Each whole-mounted tumour was subjected to tissue clearing and fluorescence microscopy to score the number of clones per tumour (Figure 3H-J). Out of 184 tumours analysed, 84 did not contain clones (45%). Dividing the data into early (day 13 post Ru486 (N=2)) and late (day 17 and 19 post Ru486 (N=2)), demonstrated that at the earlier time tumours contained more clones than later ones; 20% and 60% of tumours containing no clones respectively (Figure 3I). Moreover, there appeared to be a trend that early tumours with clones contained more clones than late tumours (Figure 3I). The average number of clones per tumour (of all tumours with clones, N=100) was found to be 4.4 and the highest number of clones in one tumour was 28 (Figure 3J). Tumour size appeared to correlate with number of clones, so that larger tumours contained a higher number of clones than smaller tumours (Figure 3K). Together these results suggest that tumour size dictates the number of clones per tumour that can be induced by Dre^Pr^ and that as predicted by a neutral competition model of stem cell replacement there are clonal extinction events occurring within developing tumours with time.

Next we sought to exploit the different times that mice were culled post lineage induction to derive the neutral behaviours of clones in tumour glands and to compare these to that of normal epithelium and of the background tumour-prone epithelium. Clone sizes were quantified by classifying each clone by the proportion of the gland it occupied (as a fraction of 8) within and outside tumours and their spatial distribution determined with respect to the tumour centre (Tc) (Figure 4A) (see Methods). Analysis within all tumours showed no clear relationship between clone size and tumour size (Figure 4B). However, although changes in tumour clone size distributions between 13 and 19 days were roughly parallel to that predicted for wildtype tissue and the background tissue in the same animals there was a massive over representation of monoclonal glands within tumours that did not fit with the trajectory of these later time points (Figure 4C). Specifically, at 13, 17 and 19 days post-induction, 25.3% (69 of 273), 28.4% (23 of 81) and 37.2% (32 of 86) of surviving clones inside tumours had monoclonally converted while outside tumours 3.1% (4 of 130), 7.0% (5 of 71) and 12.5% (8 of 64) of surviving clones were monoclonal. Wildtype neutral drift theory in the small intestine predicts 2.2%, 5.5% and 7.8% at these three timepoints (Figure 4C). Together, this suggests that either the clone dynamics of tumours changes over time or that they are heterogeneous with some glands showing accelerated dynamics. The spatial distribution of clones and tumour size were explored to determine if different behaviours related to proximity to the tumour edge or the size of the tumour, but no obvious trend was observed (Figures 4D and S5A). To explore whether this effect was due to some glands showing accelerated dynamics (i.e the monoclonal glands) the monoclonal proportion in the tumour dataset was rescaled to be identical to the data recorded outside of tumours. This analysis showed that even after rescaling, the intracrypt clonal dynamics within tumours was accelerated compared to that of external crypts (Figure 4E). Such variable behaviour precludes deriving stem cell metrics using neutral drift theory but indicate that the tumour glands arising from stabilisation of β-catenin all show accelerated but variable clone dynamics leading to monoclonality.

**Figure 4:**
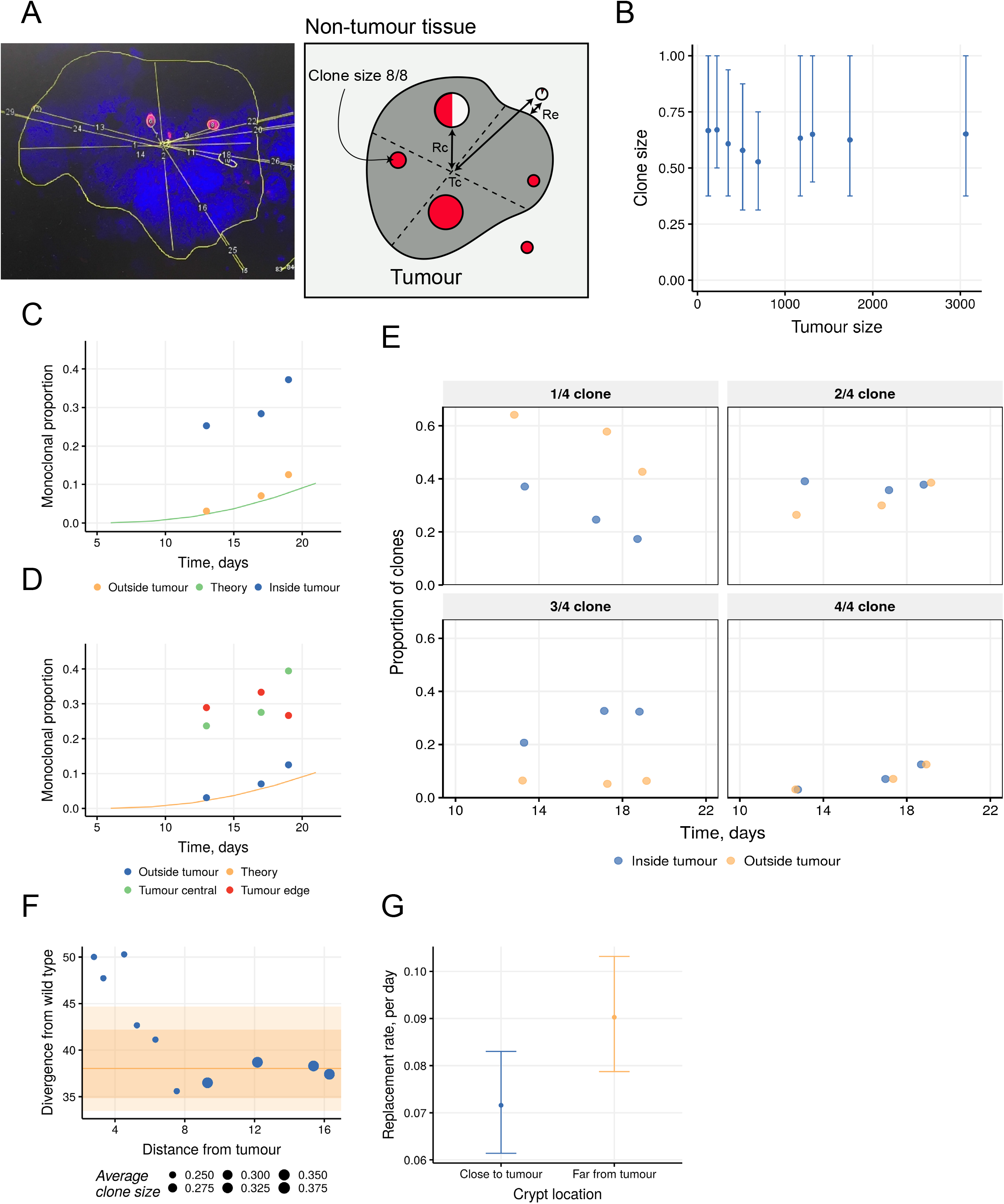
Stem cell dynamics are accelerated within *Ctnnb1* driven tumours but are inhibited adjacently to them. **(A)** Schematic diagram and real example of measurements used to characterise clones inside and outside of tumours. **(B)** Graph displaying average clone size across binned median of tumour sizes. Each bin contains 50 clones. Points and bars show median and interquartile range. **(C)** Graph comparing the proportion of surviving clones that have monoclonally converged within tumours and in surrounding tissue. **(D)** As in (C) but with the intra-tumour data split into those clones that are further from (tumour edge) or closer to the tumour centre (falling without or within 75% of the average tumour radius respectively). **(E)** The proportion of surviving non-fixed clones (in crypt quartiles) inside and outside tumours evolve differently in time when the monoclonal proportion inside tumours has been rescaled to be identical to the monoclonal proportion outside tumours. **(F)** Divergence from theoretical wild type of the clonal dynamics in crypts occupying adjacent intestinal epithelium surrounding ß-catenin tumours. Clones are grouped into overlapping bins containing equal numbers of crypts (100 clones per bin) increasing in distance from the edge of the tumour. Line and ribbon shading show the 80% and 95% credible intervals around the median of 1000 simulated data sets of 100 wildtype crypts. Point sizes show the average clone size in each bin. Distances are measured in crypt diameters. (**G)** The daily replacement rate λ of stem cells undergoing neutral drift is inferred for clones in tumour-adjacent epithelium. Close to tumour (Re < 7 crypt diameters) λ = 0.07 (80% confidence interval [CI]: 0.061–0.083); far from tumour (Re ≥ 7 crypt diameters) λ = 0.09 (80% CI: 0.079–0.103).

In considering the clone dynamics of the background clones in tumour bearing mice we next considered the impact of tumours on their behaviour. The size distribution of clones was determined with respect of their proximity to tumours by applying a rolling window moving away from the tumour that always contained the same number of clones. This analysis revealed that crypts closer to tumours had appreciably smaller clones reflecting slower drift dynamics and longer times to achieve monoclonality despite not being appreciably different in size (Figures 4F and S5B). This trend did not vary with the size of tumours (Figure S5C). Calculation of stem cell replacement rates confirmed that these were reduced in tumour adjacent crypts (Figures 4G and S5D). These findings suggest that tumours create an inhibitory local environment that slows stem cell replacement processes in adjacent crypts.

Together these observations show that RDre^Pr^ can be used for lineage tracing to reveal the altered stem cell behaviours arising as a consequence of stabilisation of β-catenin and in the resultant tumours.

## Discussion

The bulk of somatically induced, genetically engineered mouse models depend on the regulated activity of Cre-recombinase. Use of appropriate promoter elements and/or post-translational ligand-dependent regulation allows gene specific changes to be mediated at different developmental stages, in specified tissues, at known times and to varying extents. In parallel, Cre-activated reporters have offered a route to determining altered cell fates and properties. Here we have shown that Dre^Pr^ is activated in a dose-dependent manner in various tissues and can be used for lineage tracing in the small intestine and colon with a robustness that allows for detailed interpretation of quantitative data. In addition, Dre^Pr^ can be combined with Cre-mediated models as shown by the lineage tracing following stabilisation of ß-catenin driven tumours.

A prerequisite for lineage tracing is to have minimal reporter background recombination in the absence of inducer. Importantly, RDre^Pr^ mice show no background activity. In contrast, models that rely on highly efficient recombination to achieve tissue wide alterations in gene expression (e.g. *Villin*^CreErt^ in intestinal studies) are often subject to significant background recombination. It follows that such models are inappropriate for sporadic induced recombination or clonal lineage tracing. Hence the RDre^Pr^ mouse line is an appropriate tool for low frequency recombination with minimal background.

This is the first description of a recombination driven sequential model for lineage tracing in intestinal tumours. The results show that clone bearing tumours in the AhCre^Ert^;*Ctnnb1*^lox(ex3)^;RDre^Pr^;RtdTom^rsr^ model contains an average of 4 clones at the end of the experimental animal life span. The surviving clones observed in the 2-3 week interval following lineage tracing have predominantly populated whole tumour glands. In this regard the current findings are consistent with our previous study using an approach of ‘continuous labelling’ in which analysis of spontaneous mutations within the glands of spontaneous *Apc*^min^ adenomas indicated that these are maintained by a small number of clonogenic stem cells between which there is a high rate of replacement. However, here using the additional resolution of a pulse-chase approach mediated by Dre^Pr^ we further identify heterogeneity in the clone dynamics of individual tumour glands in the *Ctnnb1*^*l*ox(ex3)^ model.

In considering the clonal dynamics of the background epithelium in intestines heavily ‘peppered’ with tumours we determined that proximity to a tumour slows the inferred rate of stem cell replacement. Many reports have indicated that the impact of the mutations that cause colorectal cancers by hyperactivating the Wnt signalling pathway also cause induction of secreted negative feedback inhibitors or of other pathways that would normally act to downregulate Wnt signalling (González-Sancho et al., 2005; Kakugawa et al., 2015; Koo et al., 2012; Mishra et al., 2019). It appears likely that some of these are acting in a non-cell (or gland) autonomous manner and are additionally slowing the stem cell competitive replacement process in nearby crypts. Determining the significance of this phenomenon is beyond the scope of this report but we speculate that there may be a form of inter-gland competition such that tumour glands promote their growth by reducing the fitness of adjacent wildtype crypts.

Together, these observations demonstrate the utility of mammalian expressed Dre^Pr^ specifically and of secondary lineage tracing in general to describe and understand complex pathologies that progress with time. The approach may be particularly relevant to documenting the nature and efficacy of therapeutic interventions applied at different stages of disease progression.

## Acknowledgments

This work was supported by Cancer Research UK (to A.K.T, R.K., M.C, F.L and D.J.W.) and by the Wellcome Trust, grant 103805 (to A.K.T and D.J.W). We thank the Histopathology and Pre-Clinical Genome Editing cores and the Biological Resources Unit at the CRUK Cambridge Institute for transgenic generation and technical support.

## Author Contributions

Conceptualization, D.J.W., A.K.T., and R.K.; Methodology, R.K., D.J.W., and A.K.T.; Formal Analysis, A.K.T, D.K., and E.M.; Investigation, A.K.T., D.K., M.C and F.C.L; Writing – Original Draft, D.J.W., and A.K.T.; Writing – Review & Editing, A.K.T., D.K,, D.J.W., and E.M..; Visualization, A.K.T.; Supervision, D.J.W., R.K., and E.M.; Project Administration, D.J.W.; Funding Acquisition, D.J.W.

## Methods

### Treatment of animals

The mice were housed under controlled conditions (temperature (21 ± 2°C), humidity (55 ± 10%), 12h light/dark cycle) in a specific-pathogen-free (SPF) facility (tested according to the recommendations for health monitoring by the Federation of European Laboratory Animal Science Associations). The animals had unrestricted access to food and water, were not involved in any previous procedures and were test naive. Mice used in this study were 8-16 weeks old males and females of C57BL/6 background. <u>Ru486</u> (Sigma, cat. M8046) was dissolved in 100% ethyl alcohol (Honeywell, cat. E7023) to make a 50 mg/mL solution and further diluted into a working solution of 10 mg/mL in 40% cyclodextrin/H_2_O. <u>ß-naphthoflavone</u> (Sigma, cat. N3633) was dissolved to a working solution of 8 mg/mL in corn oil (Sigma, C8267). <u>Tamoxifen</u> (Sigma, cat. T5648) was dissolved in 100% ethyl alcohol (Honeywell, cat. E7023) to make a 200 mg/mL solution and further diluted to a working solution of 20 mg/mL, in sunflower oil (Sigma S5007).

### Subcutaneous insertion of Ru486 slow release pellet(s) in mice

This procedure was carried out by the Biological Resource Unit Core at CRUK CI. Mice receiving pellet insertion surgery were all 8-10 weeks of age. Mice were placed under general anaesthesia and a 1 cm subcutaneous incision was made on the back flank of each mouse in which 1-3 Ru486 pellets (Innovative Research of America, 10 mg/pellet, 90 days release, cat. NX-999) were placed. The wound was closed with surgical glue.

### Model creation

The RtdTom^rsrlsl^ mouse was bought from the Jackson Laboratories (stock no. 021876). The deletion of the lsl cassette to generate the RtdTom^rsr^ strain was carried out by the *CRUK CI Genome Editing Core*. RtdTom^rsrlsl^ early embryos were cultured with the cell permeable TAT-Cre *in vitro* and embryo transfer was performed as described by (Ryder et al., 2014). PCR screening was used to ensure lsl cassette deletion. <u>RDre and RDre^Pr^:</u> were generated in house at the *CRUK CI Genome Editing Core* in the Biological Resource Unit by embryonic stem cell electroporation and homologous recombination of *R26* targeting vectors and subsequent oocyte injections. A splice acceptor site was inserted immediately before the Dre sequence and Pr (consisting of the Ru486 responsive mutant hormone binding domain of the human progesterone receptor hPR891, Kellendonk et al., 1996) was fused to the N-terminus of Dre (fusion site identical to Anastassiadis *et al*., 2009) followed by a Bovine Growth Hormone (BGH) poly-adenylation signal and inserted into a R26 targeting vector (Vooijs, et al., 2001). The RDre model lacks the Pr sequence. ES cell screening for correct 5’ integration was carried out by PCR amplifying with P1 (AGAGTCCTG-ACCCAGGGAAGACATT) and P2 primers (CATCAAGGAAACCCTGGACTACT-GCG). 3’ end integration was confirmed with primers P3 (GTCACCGAGCTGCAA-GAACTCT) and R2 (GGTGGTGGTGGTGGCATATAC-ATT). Single copy insertion of the vectors was confirmed by copy number assay before oocyte injection of ES cells.

### Animal genotyping

Genotyping was carried out by Transnetyx Inc.

### Epithelial cell isolation

Small intestine and colon were dissected, flushed with PBS, everted and fed onto a glass rod spiral. They were incubated at 37°C in Hank’s Balanced Salt Solution (HBSS) without Ca^+2^ and Mg^+2^, containing 10 μM EDTA and 10 mM NaOH. Epithelial cell release was facilitated using a vibrating stirrer (Chemap). Samples were incubated for 1 h and pulsed every 10 min. Fractions were collected after each pulse, and fresh solution added. Fractions were pooled and washed in cold 2% FBS/PBS. Samples were snap frozen before DNA, RNA or protein isolation, or stained with antibodies for flow cytometry analysis.

### Flow cytometry

Single cell suspension obtained by trypsin treatment was washed and incubated with an anti-mouse CD326 (EpCAM) AlexaFluor 647 antibody (1:2000, clone G8.8, Biolegend). DAPI (10 μg/mL) was added to distinguish between live and dead cells. Flow sorting was carried out on a BD FACS Aria SORP (BD Biosciences), using appropriate single-stained and unstained controls.

### Whole-mounting

Tissue was cut open, pinned out luminal side up, and fixed for 3 h at room temperature in ice-cold 4% PFA in PBS (pH 7.4). Whole-mounts were washed with PBS, and incubated with demucifying solution (3 mg/mL dithiothreitol (DTT), 20% ethanol, 10% glycerol, 0.6% NaCl, 10 mM Tris, pH 8.2) for 20 min, and mucus removed by washing with PBS.

### Lineage tracing and clone quantification

For lineage tracing mice were induced with a pulse of ß-naphthoflavone and tamoxifen (40 mg/kg and 0.15 mg, respectively) or Ru486 (50 mg/kg) and tissue was whole mounted at day 4, 7, 10, 14 and 21 after pulse administration. Clone sizes were scored manually under a fluorescent microscope using the 550 nm filter. 2 cm of tissue was mounted muscle side up, on a glass-slide and clone sizes were scored as fractions of 8. All tissue was scored blinded. 3 mice per timepoint were quantified. In RDre^Pr^;RtdTom^rsr^ animals an average of 100-250 and 85-170 clones/animal/timepoint were quantified in the small intestine and colon, respectively. In AhCre^Ert^;RT^lsl^ animals an average of 200-300 and 175-300 clones/animal/timepoint were quantified in the small intestine and colon, respectively. Highest clone numbers were found at earliest timepoints. For number of clones/cm quantification, the number of clones was scored in 2-3cm of whole-mounted tissue as described above.

### Tissue clearing of tumour tissue

Whole-mounted tumour tissue was fixed in 4% PFA overnight at 4°C. Hereafter, tissue was washed in PBS for 8-hours at room temperature. Tissue was then incubated in CUBIC-1A solution (10% Triton, 5% NNNN-tetrakis (2-HP) ethylenediamine (Sigma, 122262), 10% Urea, 25mM NaCl) with DAPI 1:100 (10 mg/mL stock) at 37°C 60 RPM for a total of 5 days. On day 2 and 4 CUBIC-1A + DAPI was refreshed (Susaki and Ueda, 2016). After CUBIC-1A incubation, tissue was washed in PBS for 2 hours and then placed in RapiClear (SunJin Lab., cat. RC152002) and incubated at room temperature until tissue was see-through. Hereafter, tissue was mounted on 1 mm inserts (iSpacer, SunJin Lab.) on glass-slides in RapiClear and subjected to imaging on a TCS SP5 confocal microscope (Leica).

### Antibody staining of whole organs

<u>Whole-mounted intestinal tissue:</u> sections were washed in 0.1% PBS-T for 2 days, and blocked in 10% donkey serum in PBS overnight at 4°C, incubated with an anti-mouse CD326 (EpCAM) AlexaFluor 647 antibody (1:100, clone G8.8, Biolegend 118201), Ulex-Lectin 488 (1:100, Sigma 19337) and DAPI (10 μg/mL). Finally, the tissue was washed with PBS-T for 1 day before imaging. <u>Whole-mounted intestinal tissue carrying tumours</u> was covered in OCT and placed at −80°C over-night, then washed in 0.5% PBS-T for 2 days at 4°C and blocked in 10% donkey serum over-night. The tissue was then stained with ß-catenin (1:100, Cell Signalling 9587) and DAPI (10 μg/mL) in PBS-T for 3 days at 4°C, washed for 1 day, incubated with donkey anti rabbit 488 secondary antibody (1:500, Thermo Fisher, A-21206) for 2 days at 4°C, followed by a 1-day wash in PBS-T. Hereafter, the tissue was placed in Rapi Clear (SunJin Lab., cat. RC152002) and incubated at room temperature for 2-days before imaging. <u>The bladder, trachea, oesophagus and caecum:</u> were whole-mounted and then incubated in CUBIC1-A for 5 days (see tissue clearing), washed in 0.5% PBS-T for 2 days at 4°C, blocked in 10% donkey serum in PBS overnight and then incubated with Anti-pan Cytokeratin (1:100, Abcam ab236323) and DAPI (10 μg/mL) for 3 days at 4°C. The tissue was washed and incubated with donkey anti rabbit 488 secondary antibody (1:500, Thermo Fisher, A-21206) for 2 days at 4°C, followed by another 1-day wash in PBS-T before being placed in Rapi Clear (SunJin Lab., cat. RC152002) and incubated at room temperature until see-through.

### Immunofluorescence

tissue was excised and fixed for 48 h in 4% PFA in PBS at 4°C, after which it was transferred to 20% sucrose solution. After cryosectioning antigen retrieval was accomplished by incubating the slides in 1% SDS for 5 min. Blocking was performed with 10% donkey serum. Following wash, primary antibodies were added and incubated overnight at 4°C. The following primary antibodies were used: rabbit FITC-anti-Lyz (1:400, Dako, F037201), rabbit anti-Muc2 (1:50, Santa Cruz, sc-15334), rabbit anti-ChgA (1:100, Abcam, ab15160), and rabbit anti-Dclk1 antibody (1:1000, Abcam, ab31704). Secondary detection was with AlexaFluor 488 donkey anti-rabbit secondary antibody (1:500, Thermo Fisher, A-21206) and DAPI (10 μg/mL). Fluorescent imaging was carried out on a TCS SP5 confocal microscope (Leica).

### Immunohistochemistry

The small intestine and colon were opened and fixed for 24 hour in 4% PFA. The tissue was paraffin embedded and sectioned. RFP and ß-catenin immunohistochemistry were carried out using a Bond Max autostainer (Leica), with sodium citrate, pH 6.0 (10mM) antigen retrieval. Slides were blocked with 3% hydrogen peroxide, followed by incubation an Avidin/Biotin Blocking Kit (Vector Laboratories). Anti-RFP (1:100, Abcam ab34771) and ß-catenin (BD biosciences 610154, 0.25 ug/mL) primary antibodies were used. For ß-catenin IHC a mouse-on-mouse blocking step was added (Vector, MKB-2213). Secondary antibodies were; biotinylated donkey and biotinylated donkey anti-rabbit (1:250, Jackson ImmunoResearch, 711-065-152) and biotinylated rabbit anti-mouse IgG1 (1:500, Abcam ab125913). Slides were incubated with Streptavidin coupled with horseradish peroxidase (HRP), and colour developed using diaminobenzidine (DAB) and DAB Enhancer (Leica).

### Clonal analysis in tumours

to quantify clone sizes in and outside tumours as well as their spatial distribution, we defined each tumour centre (Tc), distance from clones inside to Tc (Rc), distance from clones in adjacent tissue to tumour edge (Re), clone sizes (as a fraction of 8) as well as crypt sizes of clones in each tiled image from animals presented in Figure 3C-K. In 4 animals a total of 100 (25 per animal) tumours were analysed.

### Computational analysis

#### Neutral drift model

As described in previous work (Lopez-Garcia et al., 2010; Snippert et. al. 2010; Vermeulen et al., 2013; Kozar et al., 2013) the intra-crypt clonal dynamics of stem cells in the murine small intestine and colon can be accurately described via the stochastic neutral drift theory, wherein a subset N of equipotent crypt base columnar cells undergo a continuous process of replacing their neighbours or themselves being replaced, with replacements occurring at a daily rate λ. The time evolution of this stochastic clonal expansion and contraction assuming a single stem cell is labelled at time t=0 can be captured via solution of the continuous-time Master equation for a one-dimensional random walk with absorbing boundaries at 0 and N (clonal extinction and monoclonal convergence, respectively):

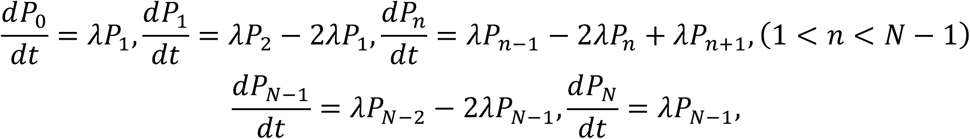

where P_n_(t) is the probability that a clone has reached a size n by time t, and P_1_(0) = 1, P_n≠1_(0) = 0 define the initial conditions. The solution of the above system (as described in Lopez-Garcia et al., 2010) is

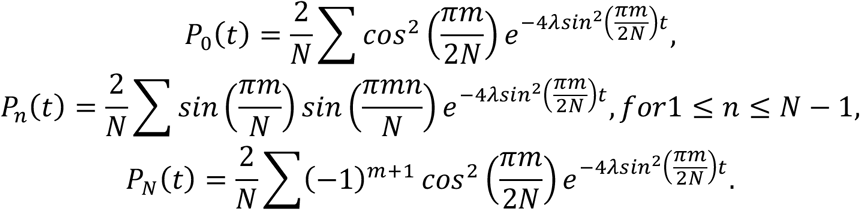

In order to fit the neutral drift model to count data of clone sizes, there are three adjustments that need to be made: (i) The probabilities P_n_(t) must be rescaled to model surviving clones, P’_n_ = P_n_ / (1-P_0_), such that the P’_n_ sum to one for n>0 for all t; (ii) The P’_n_ must be redistributed into the number of bins that were used to score the clone sizes, in this case eighths; (iii) The delay between tamoxifen administration and the accrual of the stem cell label is included as a time delay parameter τ. Then, count data X is modelled as a multinomial with P’_n_ – now a function of the model parameters λ, N and τ – as the case probabilities:

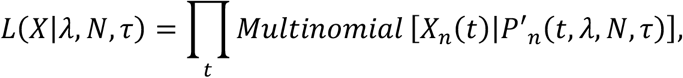

where X_n_(t) is the vector of counts of clone sizes 1≤n≤N observed at time t. The priors and associated hyperparameters were

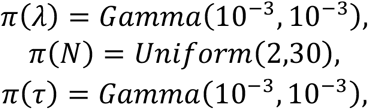

which were chosen to be uninformative.

Using this Bayesian inference model, Markov Chain Monte Carlo (MCMC) simulations were used to produce draws of the neutral drift parameters for the wildtype small intestine as shown in Fig 2K. In all MCMC sampling herein, 40000 iterations were performed on two parallel chains each with a burn in of 5000 iterations, with the results thinned by a factor of 20. From this inference, only a well-resolved value for the time delay parameter was desired to use in the tumour analysis. The time points were 4, 7, 10, 14 and 21 days post induction at which 692, 595, 592, 457 and 302 clones were observed, respectively. The parameter value was found to be τ = 2.45 (95% credible interval [CI]: 2.07–2.75). This was taken as the Dre technology-intrinsic time delay and τ_glob_ = τ was defined for use in all future computations.

#### Divergence from wild type

All tumour lineage tracing data (internal and external tissue) was taken from SI1. Therefore, the pulse-chase neutral drift theory with N_WT_ = 5 stem cells with λ_WT_ = 0.1 daily replacements (Kozar et al., 2013) was used as the baseline wild type behaviour and investigate whether and by how much the observed data diverges from it. The theoretical results were split symmetrically into 8 bins to match the eighths used to measure the clone sizes, though symmetry is broken to maintain the identity of monoclonals. This gives a theoretical distribution of clone sizes P’_n_(t), 1≤n≤8, occupied by surviving clones at a given time t post labelling, as defined in the neutral drift model section.

To investigate the effect on clonal dynamics of proximity to a tumour, the putatively normal tissue surrounding tumours was binned radially from the edge of the tumour outwards, increasing in distance from the tumour (like the rings of a target). To achieve statistical power and allow between-bin comparison these bins were allowed to overlap such that each bin contained 100 clones (thus each clone was assigned to one or more bins) but the median distance of crypts from the tumour edge in subsequent bins increased monotonically. The likelihood under the theoretical multinomial model for the pooled clone sizes in bin b and time point t, C_b_(t) is given by

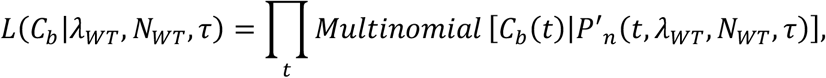

where is the number of crypts in bin b that arose from time point t. The value of the time delay τ is fixed at the value inferred from the control data as described in the previous section. The likelihood for bin b explains how well the theoretical wild type model describes the data observed in that spatial bin in tissue external to a tumour.

In order to interpret these results a null distribution on the likelihood was created by simulating counts from the theoretical multinomial model, C_sim_ ~ Multinom(P’_n_(t), n_b_), where nb is the bin size (100 clones) and where the proportion of clones from each time point was the same as that in the data. Counts were generated one thousand times and the likelihood of each simulation under the theoretical model was calculated. The credible intervals resulting from these simulations were used to judge whether the clones in the binned data were undergoing perturbed dynamics or were within the variability expected from finite sampling.

#### Quantifying the clonal dynamics close to tumours

To quantify the effect of tumours on the clonal dynamics in tumour-adjacent crypts the data set was split into two subsets: those crypts displaying dynamics not well described by the wildtype theory (those outside the 95% credible interval of the simulated null distribution, 140 crypts) and those displaying dynamics that are well described by the wildtype theory (those inside the 95% credible interval, 125 crypts). The value of the radial distance cut-off that this corresponded to is 6.99 crypt diameters (454 μm). The neutral drift replacement rate λ was inferred for each data set using a modified version of the MCMC algorithm described above, wherein the delay parameter and the number of stem cells were fixed to τ = τ_glob_ and N = N_WT_ in order to make the replacement rate identifiable.

#### Binning clones

Two methods of binning were used to group clones into statistically powered subsets with respect to a measurement, say X. First, overlapping bins with a constant number of clones in each but a variable width in the parameter X (used in Figs 4F and S5B-C, with Fig 4B having an overlap of zero bar the point at the far right which overlaps its neighbouring point in order to maintain constant bin size). Second, non-overlapping bins that equally divide the parameter X such that each bin can have a different number of clones (used in Fig S5A)

#### Rescaling monoclonals

To compare the evolution in time of partial clone sizes (those clones that have not become fixed in a gland) within tumours and in tumour-adjacent epithelium, the large monoclonal bias inside tumours was first scaled away (this is necessary as we work in proportions). The rescaled and re-normalised intra-tumour clone 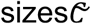 were calculated as:

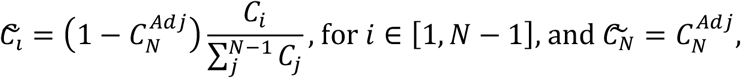

where *C* is the raw intra-tumour clone size distribution and 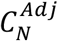 is the proportion of monoclonals in the clone size distribution of the tumour-adjacent tissue. The results are shown in Fig 4E.

