## Supplementary figure legends for "Heterogeneity in clone dynamics within and adjacent to intestinal tumours identified by Dre-mediated lineage tracing"

**Supplementary Figure 1: RDre and RDre$^{Pr}$ animals ubiquitously express Dre in tissues from different germ layers. (A)** Schematic representation of the targeted locus on *R26* following a RDre or RDre$^{Pr}$ targeting. End (endogenous *R26* locus), Hom (homology arm), SA (splice acceptor), BGH (bovine growth hormone poly-A signal), PGK (Phosphoglycerate Kinase promoter), Puro (puromycin selection cassette). **(B)** Representative IHC images (N=4 per group) of tdTom expression in various tissues from RDre;RtdTom$^{rsr}$ and RDre$^{Pr}$;RtdTom$^{rsr}$ animals either uninduced (UI) or surgically implanted with 1-3 90-day slow-release 10mg Ru486 pellets, 75 days post implantation. Pictures are 10X magnification.

**Supplementary Figure 2: not all tissues express Dre in Dre$^{Pr}$ animals following Ru486 activation.** Representative IHC images of tdTom expression in tissues from RDre;RtdTom$^{rsr}$ and RDre$^{Pr}$;RtdTom$^{rsr}$ animals either uninduced (UI) or surgically implanted with 1-3 90-day slow-release 10mg Ru486 pellets, 75 days post implantation. The lung, bone, tongue and heart of RDre$^{Pr}$ animals did not contain tdTom+ cells after Ru486 exposure.

**Supplementary Figure 3: Dre$^{Pr}$ is activated in a Ru486 dose-dependent manner in the small intestine (A)** Representative pictures of proximal small intestine and colon of uninduced (UI) and 1, 2 or 3 Ru486 slow release pellet implanted R_Dre$^{Pr}$;RT$^{rox}$ animals 75 days post implantation. Black colour represents tdTom signal (N=4 per group). **(B)** Representative flow cytometry analysis of proximal small intestine from above animals. Shown are the gating strategy for all cells, single cells, live cells and EpCam+/tdTom+ and representative experimental plots. Cells were stained with EpCam Alexa-647 and DAPI. All gates were set on an unstained and uninduced control sample. (N=4 per group) **(C)** Bar-plot summarising percentage of EpCam+/tdTom+ cells quantified by flow cytometry in (B). Each data-point represents one animal, error bar display SD.

**Supplementary Figure 4: Gating strategy for Figure 3B**. All gates were set on an un-stained uninduced control sample.

**Supplementary Figure 5: Mathematical modelling of stem cell dynamics in adjacent and intra tumour crypts (A)** Plots show absence of a relationship between clone size and tumour size. Bins contain equal numbers of clones but are of variable width along the tumour size axis. Tumour size is scaled by the median crypt area of crypts in tumour-adjacent tissue. For each time point, same-coloured points sum to one. Areas are measured in average crypt areas. **(B)** The divergence of dynamics away from wildtype in clones close to tumours is not correlated with the gland size. Clones are binned as in Fig 4F. Distances are measured in crypt diameters. **(C)** As in (B) showing that the altered dynamics close to tumours does not correlate with tumour size. **(D)** Plot shows density of the Markov Chain Monte Carlo samples of the posterior distribution for the stem cell replacement rate in crypts close to and far from tumour. Histogram showing the distribution of the replacement rate as a function of sampling density in data close to tumour and far from tumour.
