## Supplementary figures and images for "Heterogeneity in clone dynamics within and adjacent to intestinal tumours identified by Dre-mediated lineage tracing"

### Supplementary figures

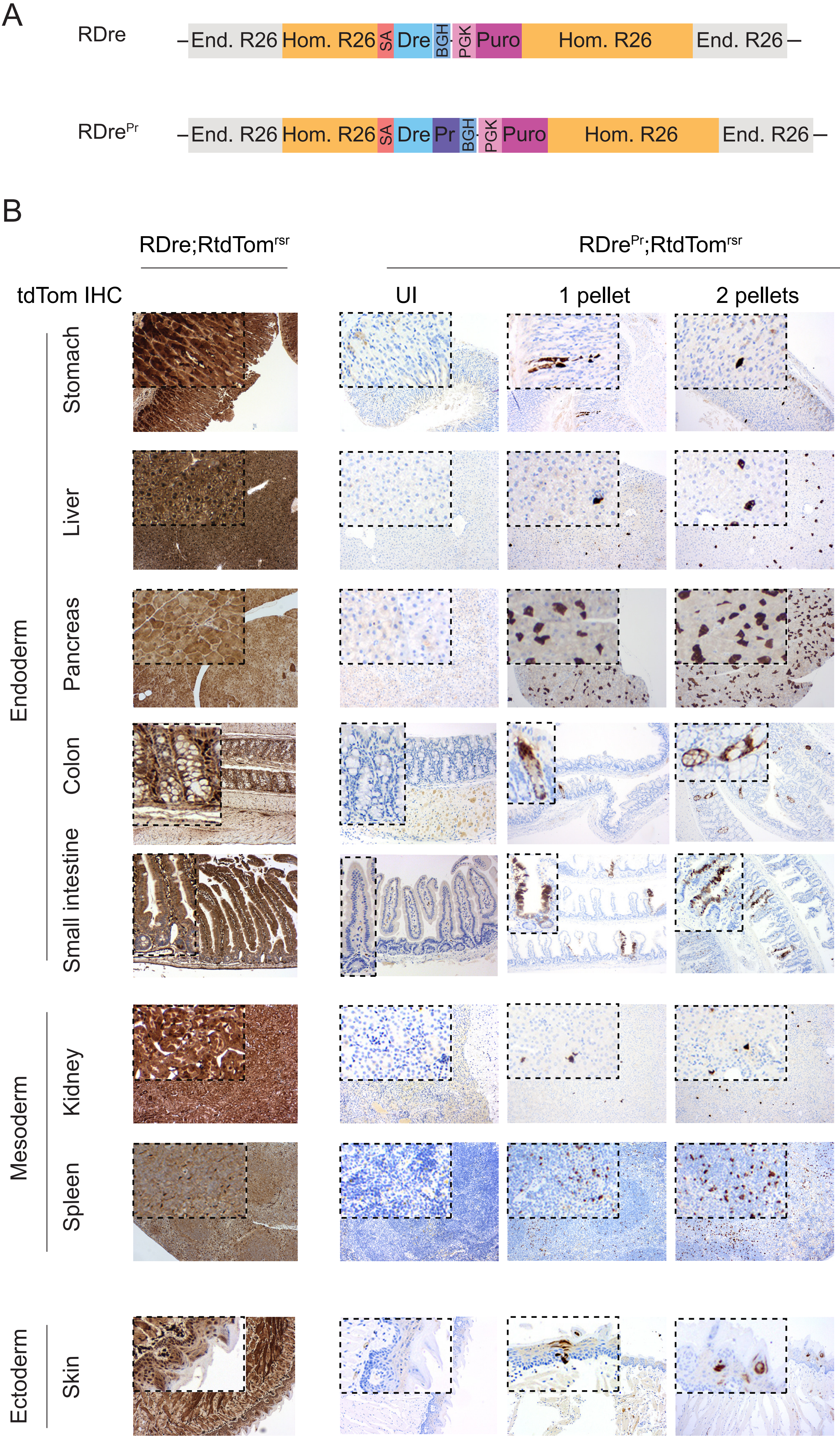

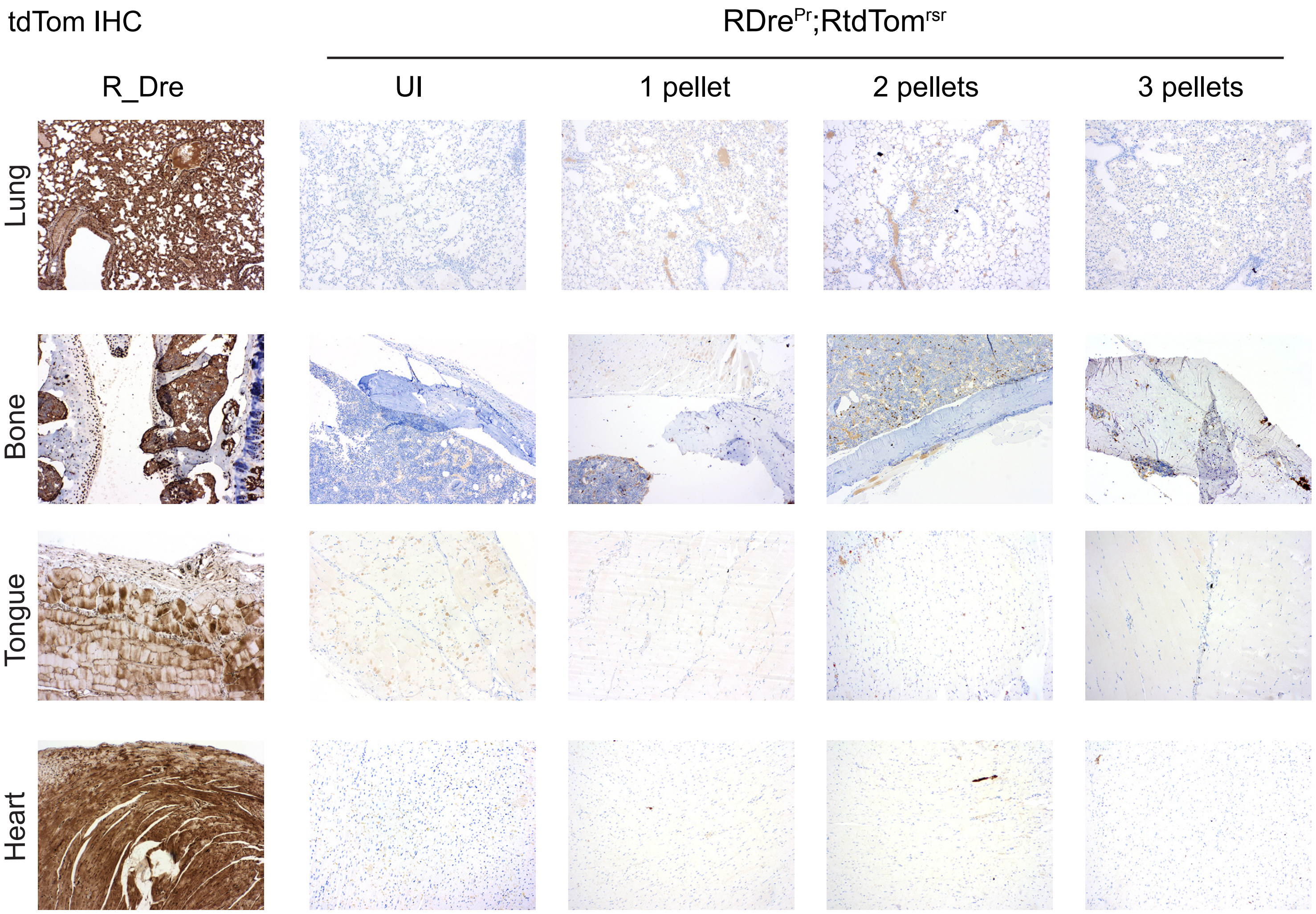

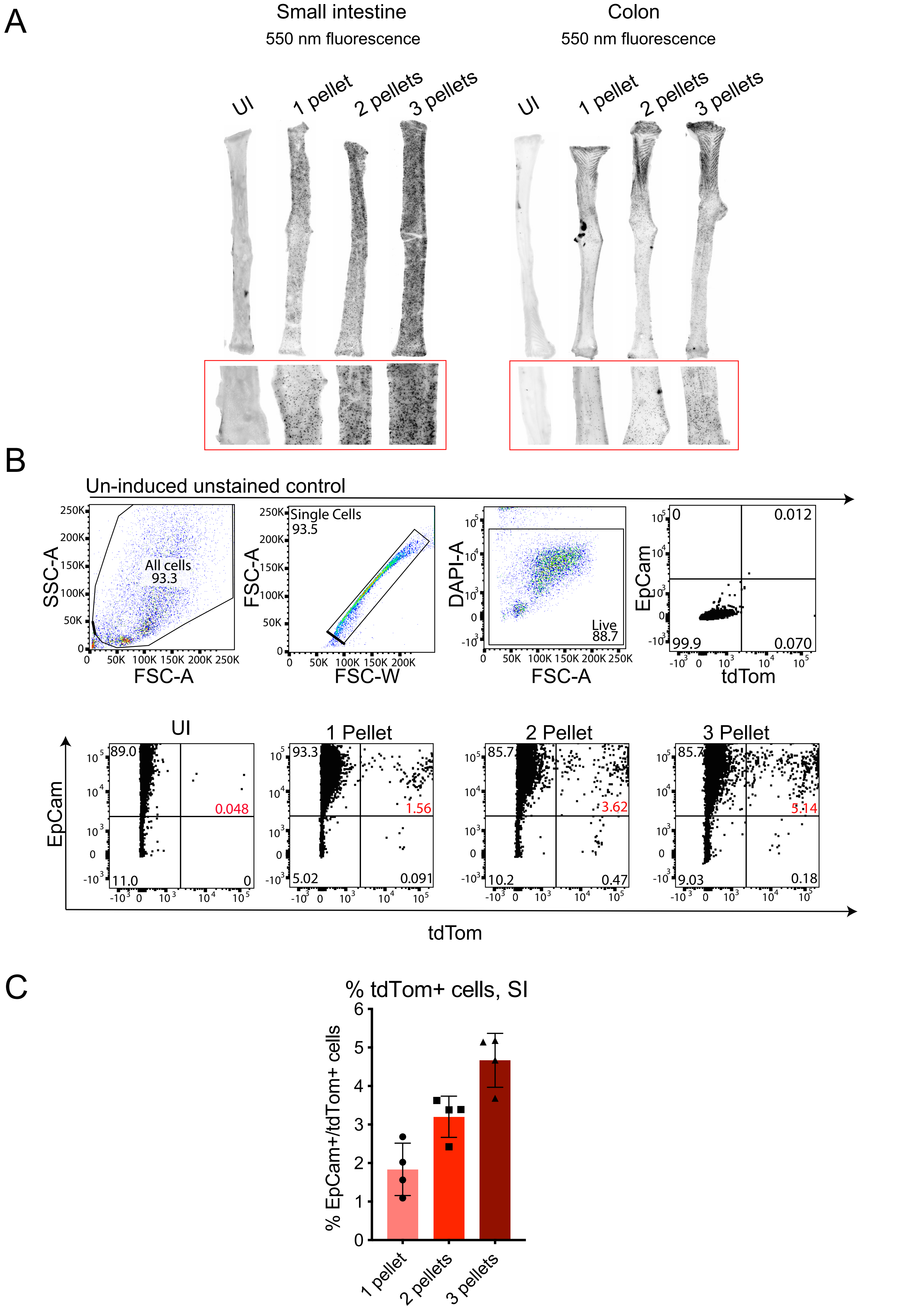

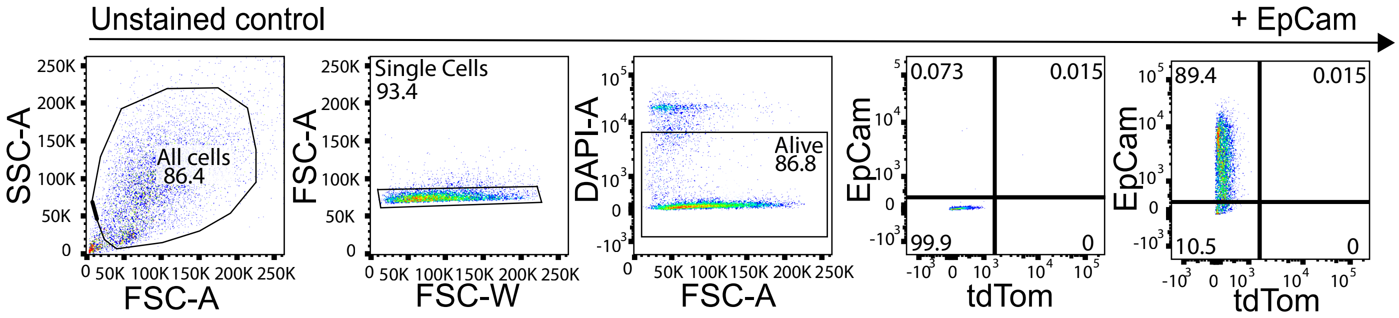

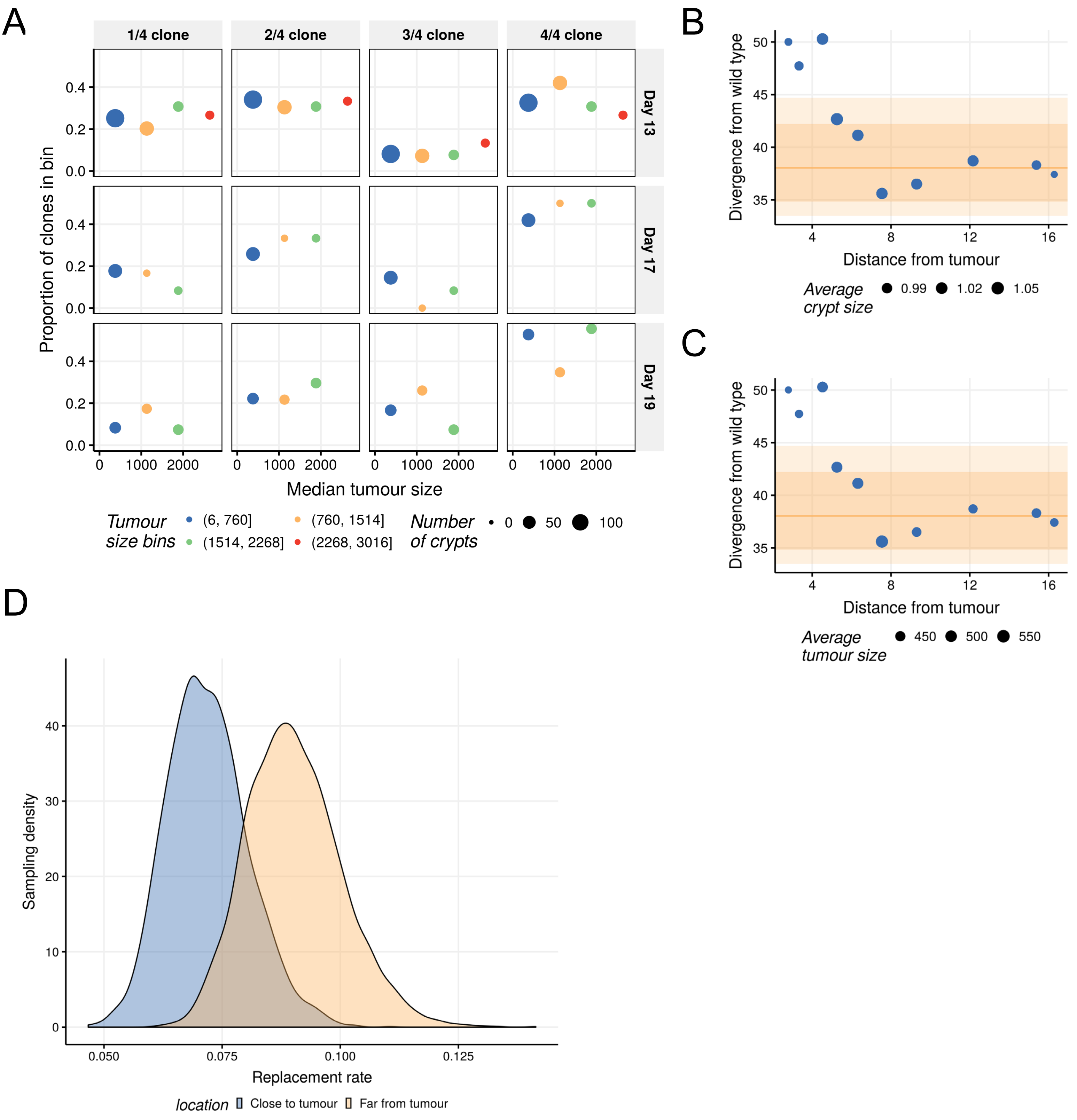
